# Novel Role of AcylCoA:cholesterol acyltransferase 1 (ACAT1/SOAT1) in Diabetic Retinopathy

**DOI:** 10.1101/2025.10.09.681484

**Authors:** Syed Adeel H. Zaidi, Tahira Lemtalsi, Zhimin Xu, Mai Yamamoto, Porsche V. Sandow, Steven E. Brooks, R. William Caldwell, Ruth B. Caldwell, Modesto A. Rojas

**Author notes:** Address of correspondence for this article, Ruth B. Caldwell, PhD, Vascular Biology Center, 1460 Laney Walker Blvd, Augusta University, Augusta, GA 30912-2500, Modesto A. Rojas, MD, Vascular Biology Center, 1460 Laney Walker Blvd, Augusta University, Augusta, GA 30912-2500.

## Abstract

Hypercholesterolemia and excessive cholesterol ester (CE) production have been linked to chronic inflammation and vascular dysfunction during cardiovascular disease. Upregulation of AcylCoA:cholesterol acyltransferase 1 (ACAT1/SOAT1), the enzyme responsible for retinal CE formation, has been implicated in pathological retinal neovascularization. Here we determine the role of this process in diabetic retinopathy (DR). Ins2^Akita^ diabetic mice were treated with the specific ACAT1/SOAT1 inhibitor K604 (10 mg/Kg, i.p.) beginning at 10 weeks for 2 weeks or 8 months for 2 months. ACAT1/SOAT1 expression and CE formation were assayed along with oxidative stress, inflammation, vascular pathology, and neuronal function. ACAT1/SOAT1 expression was also assayed in human retinas and vitrectomy specimens. Retinas from early-stage Ins2^Akita^ mice exhibited increases in CE deposition, superoxide production, and expression of ACAT1/SOAT1, LDLR, TREM1, MCSF, and VEGF along with leukostasis, vascular leakage, acellular capillary formation, retinal ganglion cell loss, and impaired visual function. Late-stage increases in CE, ACAT1/SOAT, oxidative stress, inflammation, and impaired visual function were also observed. These changes were significantly inhibited by K604 treatment. The protective effects were independent of changes in systemic glucose or body weight. Human retina and vitrectomy samples also showed increases in ACAT1/SOAT1 and CE, respectively. Specific inhibition of ACAT1/SOAT1 with K604 normalizes ACAT1/SOAT1 expression and CE formation and prevents increases in oxidative stress and inflammation and preserves retinal structure and function in both early and late stages of DR. These findings identify ACAT1/SOAT1 as a promising therapeutic target for both early intervention and later stage treatment of DR.

**One sentence summary:** Inhibiting cholesterol esterification limits retinal neurovascular injury in diabetes.

## INTRODUCTION

Diabetic retinopathy (DR) is the most frequent cause of blindness in working adults in the US. An estimated 9.6 million people in the U.S. are living with DR. Of these, 1.84 million have vision-threatening DR. This number is expected to double by 2050 (*1*). This condition is associated with neuronal and glial injury as well as vascular dysfunction (*2–6*). Laser photocoagulation is usually effective for advanced retinopathy but can impair vision and in some patients, retinopathy continues to progress. Intraocular injections of anti-VEGF antibodies have shown efficacy in reducing diabetic macular edema (*7*). However, the effectiveness of this treatment is limited in some cases and resistance to intravitreal anti-VEGF and recurrence of pathology have been reported (*8*). Also, anti-VEGF treatment does not promote tissue repair and may have adverse effects, including neuronal damage, retinal detachment, and intraretinal infection (*9*). Therefore, novel therapeutic strategies are urgently needed to limit injury and promote repair during DR.

Diabetes mellitus is a chronic metabolic disease characterized by dysfunctional insulin activity and high plasma glucose levels which lead to changes in glycogen and fatty acid metabolism followed by lipid accumulation and increased oxidative stress (*10, 11*). While results of studies examining the link between DR and lipid abnormalities are inconsistent, one study found that high levels of circulating LDL cholesterol are associated with significant risk for diabetic macular edema and retinal hard exudates (*12*). Dyslipidemia has also been linked to faster progression and worsening of DR (*13*), (*14*). Also, studies of post-mortem retinas from DR patients have shown increases in oxidized LDL associated with macrophage infiltration and severity of DR (*15*), (*16*). In peripheral vascular disease, accumulation of CE stored as cytoplasmic lipid droplets is a major characteristic of macrophage foam cells that are central to the development of atherosclerotic plaques (*17*). Based on these findings, we hypothesized a pathophysiological role for CE formation in DR.

Our previous research has identified AcylCoA:cholesterol acyltransferase 1 (ACAT1/SOAT1) as a critical mediator of pathological retinal neovascularization (RNV) in a mouse model of oxygen-induced retinopathy (OIR) (*18*). ACAT1/SOAT1 is an endoplasmic reticulum membrane-bound enzyme that uses cholesterol and long chain fatty acid as substrates to produce CE (*19*). ACAT1/SOAT1 is expressed in many cell types, including photoreceptors, macrophages, monocytes, and microglial cells (*5, 20*). Our studies in the OIR mouse model have shown increases in lipid accumulation and CE formation in areas of pathological RNV. These increases were associated with significant elevation of ACAT1/SOAT1, macrophage colony stimulating factor (MCSF), LDL receptor (LDLR), and VEGF. Treatment with a specific inhibitor of ACAT1/SOAT1, K604, blocked increases in all these proteins except VEGF. Moreover, K604 treatment also inhibited RNV and improved vascular repair (*18*). In the present study we have tested the role of ACAT1/SOAT1 activity in neurovascular injury during DR. We hypothesized that diabetes causes high levels of retinal CE formation which activates inflammatory signaling pathways and therefore promotes neurovascular damage.

## RESULTS

### ACAT1/SOAT1 is upregulated and functionally active in human diabetic retinopathy

Analysis of human DR tissues revealed a robust upregulation of ACAT1/SOAT1 expression and activity. Immunolabeling of retinal sections from DR donors showed strong ACAT1/SOAT1 immunoreactivity in the photoreceptor outer segments and retinal pigment epithelium (RPE), in contrast to weak signal in non-diabetic controls, indicating diabetes-associated enzyme upregulation (**Fig. 1A**). Consistently, vitreous samples from patients with proliferative DR exhibited elevated CE levels, confirming enhanced ACAT1/SOAT1 activity in the diabetic retina (**Fig. 1B**).

**Fig. 1.**
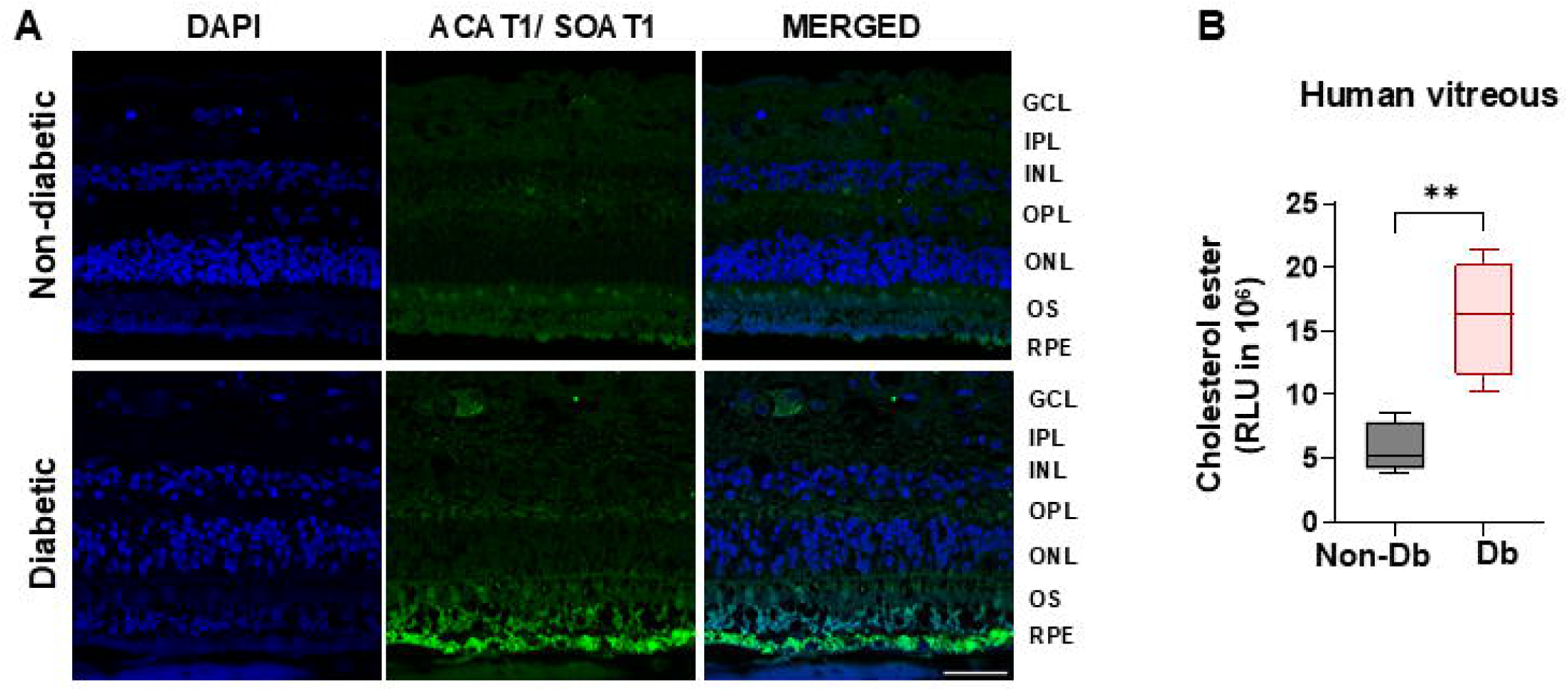
ACAT1/SOAT1 expression and activity are upregulated in retina and vitrectomy samples from patients with diabetic retinopathy (DR). **A.** Immunolabeling of retina sections from human donors with DR showed increased ACAT1/SOAT1 immunoreactivity in the photoreceptor outer segments (OS) and retinal pigment epithelial (RPE) cells, as compared with non-diabetic control donors. Donor sample information is provided in supplementary table 1. Scale bar = 50 µm. n=3. **B.** Vitrectomy tissues collected from patients with PDR showed an increased level of CE, as measured by ELISA. Donor sample information is provided in supplementary table 2. Data are presented as mean ± SEM; n=4-5; **p<0.01.

### Diabetes promotes dysregulation of lipid metabolism in the diabetic retina

To investigate the functional role of ACAT1/SOAT1 in diabetic retinopathy, we treated diabetic Ins2^Akita^ mice with the selective ACAT1/SOAT1 inhibitor K604. To validate our experimental design, we first confirmed the onset of hyperglycemia in Ins2^Akita^ mice at 5 weeks of age and established treatment regimens with the ACAT1/SOAT1 inhibitor K604 (**Supplementary Fig. 1A**). Intraperitoneal injections of K604 for 2 weeks did not affect body weight or blood glucose levels in the diabetic mice, indicating that any observed effects are independent of systemic metabolic changes (**Supplementary Fig. 1B, C**).

To determine whether diabetes alter cholesterol homeostasis in the retina, we assessed lipid and CE levels in Ins2^Akita^ diabetic mouse retinas. Oil Red O staining showed a slight increase in lipid levels in the retinal ganglion cell layer and photoreceptor outer segments of the diabetic retinas, and this was not evident after K604 treatment. Quantitation of relative fluorescence intensity (RFI) showed no significant difference in the overall RFI. (**Fig. 2A, B**). Filipin staining demonstrated prominent increases in CE deposition in multiple retinal layers of diabetic mice as compared with the non-diabetic controls and these increases were blocked by K604 treatment (**Fig. 2C, D**). Biochemical analysis further confirmed elevated CE levels in both retina (**Fig. 2E**) and plasma (**Fig. 2F**) of Ins2^Akita^ diabetic mice. These increases were blocked by K604 treatment. However, levels of cholesterol were significantly increased only in plasma, but not in retina.

**Fig. 2.**
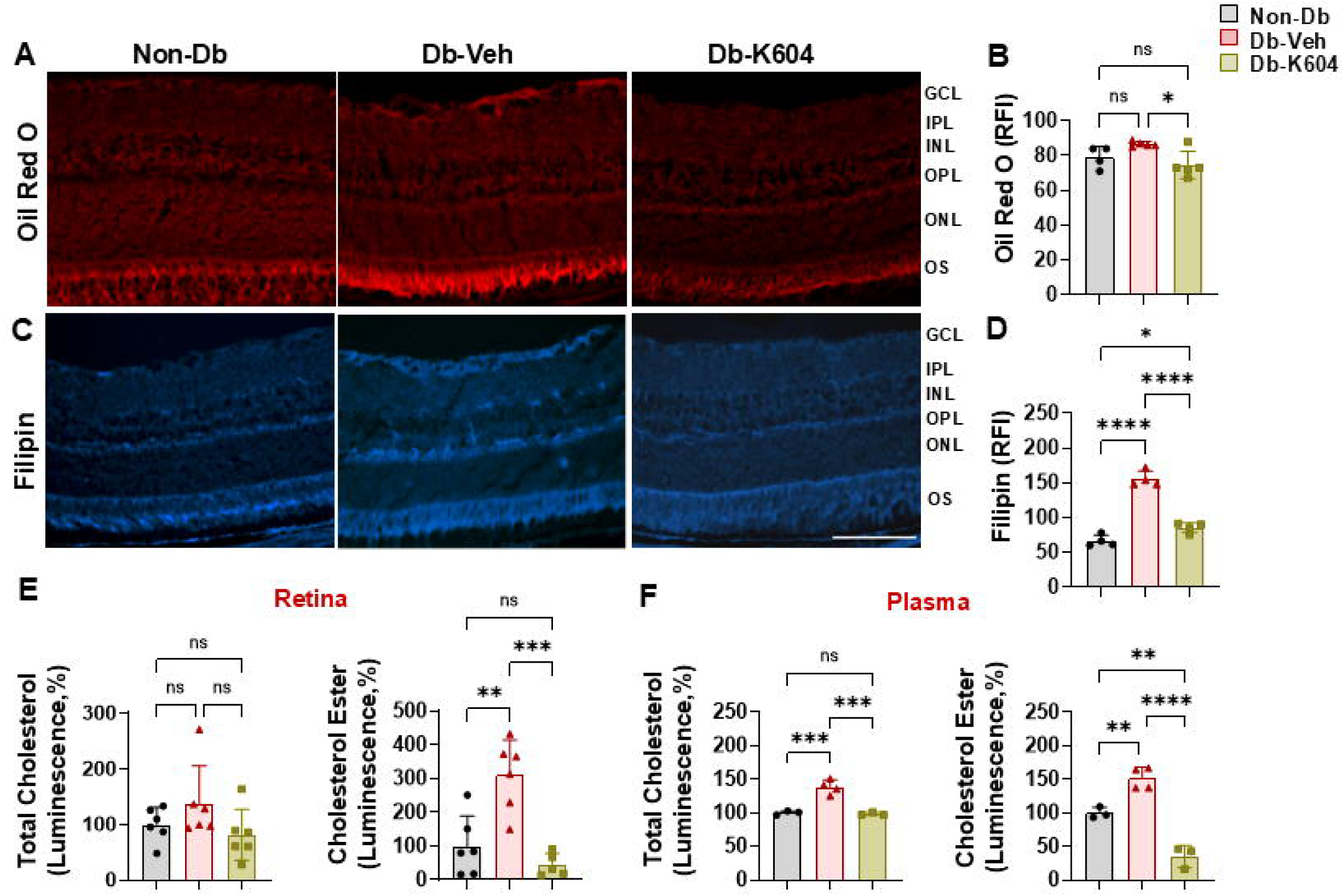
Inhibition of ACAT1/SOAT1 prevents diabetes-induced CE formation in Ins2^Akita^ mice. Ten-week old male diabetic Ins2^Akita^ mice were treated with PBS as vehicle or ACAT1/SOAT1 inhibitor, K604 (10mg/Kg, i.p.), every 2 days for 2 weeks. After 2 weeks of treatment, eyeballs and retinas were collected. **A-B.** Oil red O staining of frozen sections shows mild reactivity in the ganglion cell layer (GCL) and photoreceptor outer segments (OS) of the Ins2^Akita^ diabetic retina. Treatment with the ACAT1/SOAT1 inhibitor, K604, reduced this activity. Mean ± SEM; n=4-5; *p<0.05; ns: not significant. **C-D.** Detection of CE by filipin labeling of frozen retina sections shows prominent increases in CE formation in the ganglion cell (GCL), outer plexiform (OPL) and photoreceptor outer segment (OS) layers of the Ins2^Akita^ diabetic retina. Treatment with K604 prevented the CE increase. Scale bar = 40 µm; mean ± SEM; n=4; ****p<0.0001. **E-F.** Total cholesterol and CE levels were measured in retinas and plasma from non-diabetic control mice, Ins2^Akita^ diabetic mice, and K604-treated Ins2^Akita^ diabetic mice. This analysis showed upregulation of total cholesterol in plasma of the Ins2^Akita^ mice but not in retina. However, CE was increased in both retina and plasma of the Ins2^Akita^ mice. K604 treatment reduced retinal CE formation as well as preventing the increases in both cholesterol and CE in plasma. Data are presented as mean ± SEM; n=4-6; **p<0.01; ***p<0.001; ****p<0.0001; ns: not significant.

### ACAT1/SOAT1 inhibition normalizes expression of ACAT1/SOAT1 and suppresses inflammatory and angiogenic signaling

Western blot analyses of Ins2^Akita^ diabetic retinas demonstrated significant upregulation of LDLR and ACAT1/SOAT1 and that this was accompanied by increased expression of inflammatory and angiogenic mediators including TREM1, MCSF, VEGF, and TNFα. Treatment with K604 restored expression of all these proteins to near-control levels, suggesting that ACAT1/SOAT1 activity drives both increases of CE levels and activation of pathogenic inflammatory signaling cascades in the diabetic retina (**Fig. 3A-G**). Increased levels of ACAT1/SOAT1 in the retinal ganglion cells, outer segments, and RPE cells were also detected in retinal frozen sections from diabetic mice. These pathological increases of ACAT1/SOAT1 were blocked by the K604 inhibitor treatment, without interfering with their physiological expression (**Fig. 3H, I**).

**Fig. 3.**
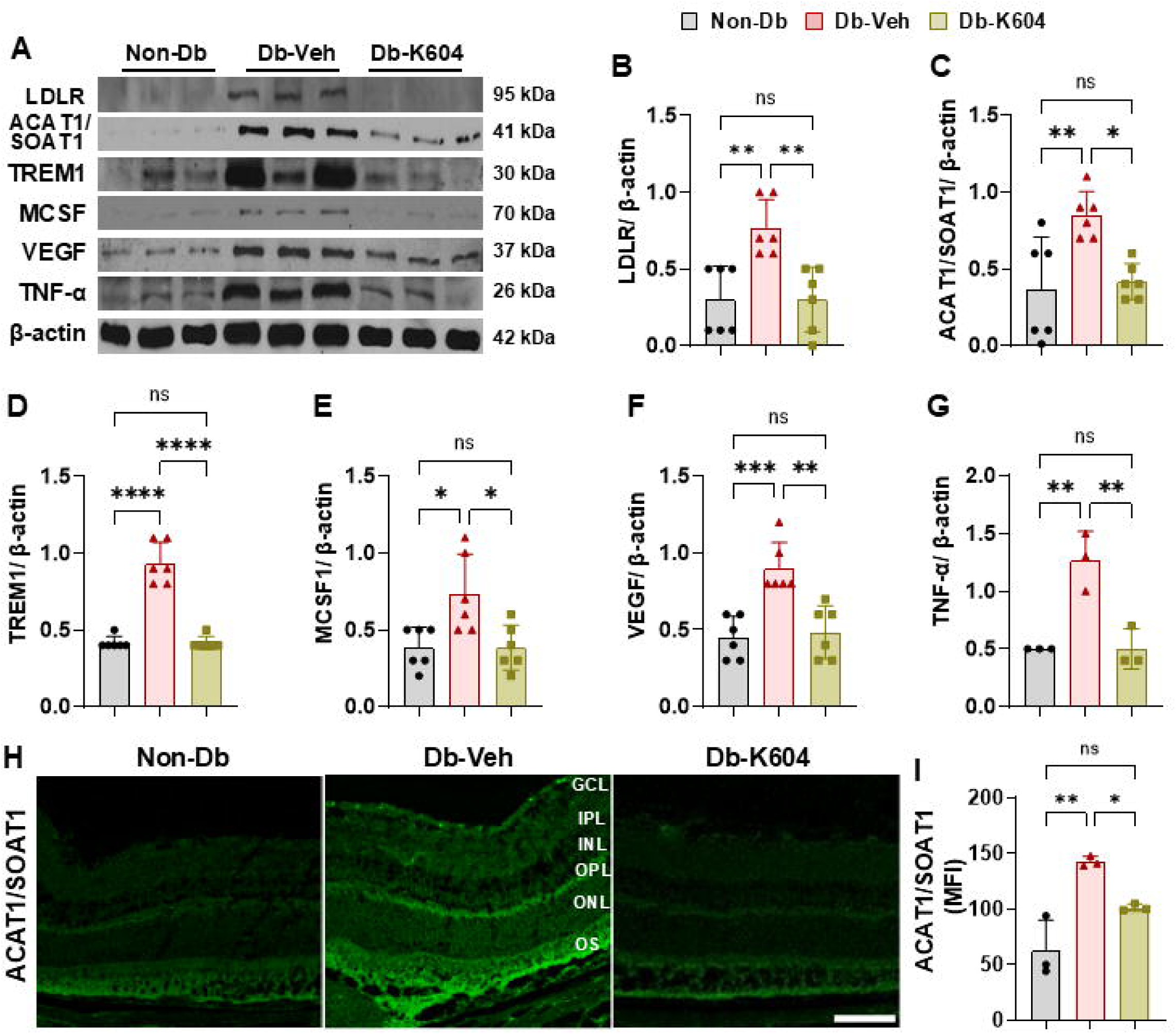
Inhibition of ACAT1/SOAT1 prevents diabetes-induced increases in expression of inflammatory mediators and ACAT1/SOAT1 in Ins2^Akita^ mice. **A.** Western blot analysis of retinas from 12-week diabetic Ins2^Akita^ mice showed marked increases in **B.** LDLR, **C.** ACAT1/SOAT1, **D.** TREM1, **E.** MCSF, **F.** VEGF, and **G.** TNF-α as compared with the WT non-diabetic controls. These increases were normalized by the treatment with K604 (10mg/Kg, ip) every two days from 10 to 12 weeks. Mean ± SEM, n=3-6; *p<0.05; **p<0.01; ***p<0.001; ****p<0.0001. **H-I**. Increased immunoreactivity for ACAT1/SOAT1 was detected in retinal frozen sections from Ins2^Akita^ diabetic mice, especially in the RGC, OPL, and OS layers. The ACAT1 immunoreactivity was significantly reduced in retinas of Ins2^Akita^ diabetic mice treated with K604. Data are presented as mean ± SEM; n=3-6; Scale bar = 25 μm.

### ACAT1/SOAT1 inhibition limits oxidative stress and retinal inflammation

Markers of oxidative stress and inflammation were elevated in diabetic retinas, changes that were mitigated by ACAT1/SOAT1 inhibition. DHE (dihydroethidium) imaging revealed that diabetes-induced superoxide production in diabetic retinas was blocked in K604-treated animals (**Fig. 4A, B**). Specificity of DHE fluorescence for superoxide was confirmed by preincubation with superoxide dismutase (SOD), which completely abolished the diabetes-induced DHE signal, (**Supplementary Fig. 2A, B**).

**Fig. 4.**
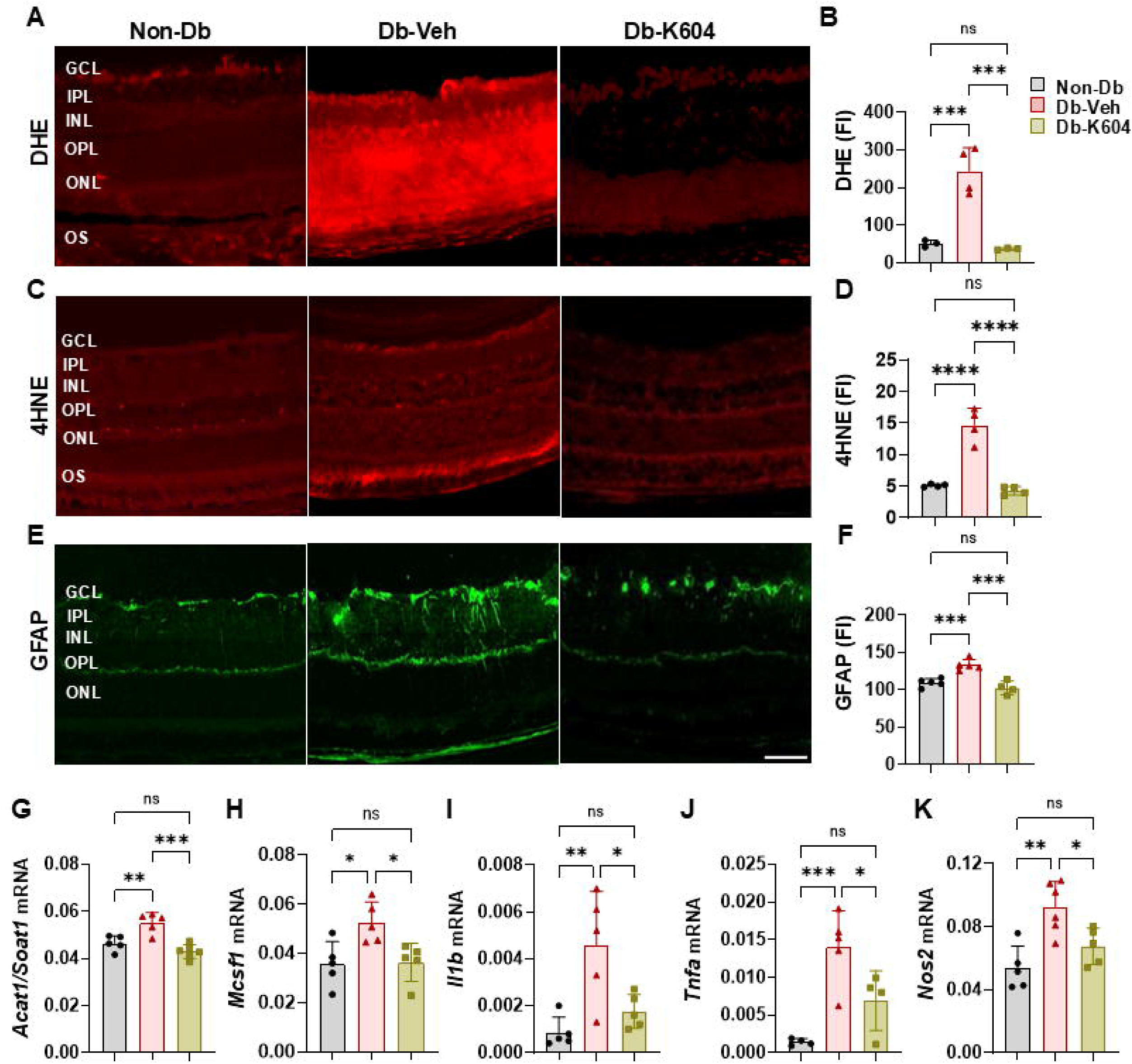
Inhibition of ACAT1/SOAT1 ameliorates the diabetes-induced increases in oxidative stress, gliosis, and expression of inflammatory mediators in Ins2^Akita^ mice. **A-B.** DHE imaging showed upregulation of superoxide formation in frozen sections from Ins2^Akita^ diabetic mice. This effect was blocked by the K604 treatment. n=4. **C-D.** Immunofluorescent labeling of frozen retina sections with 4-hydroxynonenal (4-HNE) and fluorescence intensity quantification showed increased 4-HNE levels in the ganglion cell layer (GCL), outer plexiform layer (OPL) and photoreceptor outer segments (OS) of the Ins2^Akita^ mice. K604 treatment blocked this effect. n=4. **E-F.** Immunolabeling of retina cross-sections showed increased GFAP immunoreactivity in Müller cells of the Ins2^Akita^ mice, which was blocked in the K604 treated mice. n=4. Scale bar=30 μm. **G-K.** Quantitative RT-PCR showed upregulation of *Acat1/Soat1*, along with proinflammatory mediators – *Mcsf1, Il1b, Tnfa*, and *Nos2* in Ins2^Akita^ diabetic mouse retinas, compared to non-diabetic controls. K604 treatment significantly inhibited these increases. n=4-6. All the data are presented as mean ± SEM. *p<0.05; **p<0.01; ***p<0.001; ****p<0.0001.

Labeling for the lipid peroxidation marker 4-hydroxynonenal (4-HNE) showed areas that were increased in the RGC, photoreceptors and RPE layers of diabetic mice were reduced by K604 treatment (**Fig. 4C, D**). GFAP labeling also indicated Müller glial cell activation in diabetic retinas (**Fig. 4E, F**) and this was blocked by the K604 treatment. Iba1 immunolabeling of retinal microglia revealed enlarged, ameboid microglia and macrophages in the Ins2^Akita^ diabetic retinas, but not in eyes of K604 treated animals (**Supplementary Fig. 3**). At the transcriptional level, diabetic retinas exhibited upregulation of *Acat1/Soat1* along with proinflammatory mediators including *Mcsf1*, *Il1b*, *Tnfa*, and *Nos2*, all of which were suppressed by K604 administration (**Fig. 4G-K**).

### ACAT1/SOAT1 inhibition preserves visual function in early diabetes

We next tested whether inhibiting ACAT1/SOAT1 confers neuroprotection. Confocal imaging of retinal flat mounts labeled with the neuronal cell marker RBPMS showed that K604 treatment increased the survival of retinal ganglion cells compared with vehicle-treated diabetic mice (**Fig. 5A, B**). Functional assessment using electroretinography (ERG) revealed that both scotopic a- and b-wave amplitudes were significantly improved following K604 treatment, indicating preserved photoreceptor and bipolar cell function (**Fig. 5C, D**). Moreover, visual acuity, measured by optomotor responses (OMR), was markedly enhanced in K604-treated diabetic mice (**Fig. 5E**). These findings demonstrate that the inhibition of ACAT1/SOAT1 preserves neuronal integrity and visual function in the diabetic retina.

**Fig. 5.**
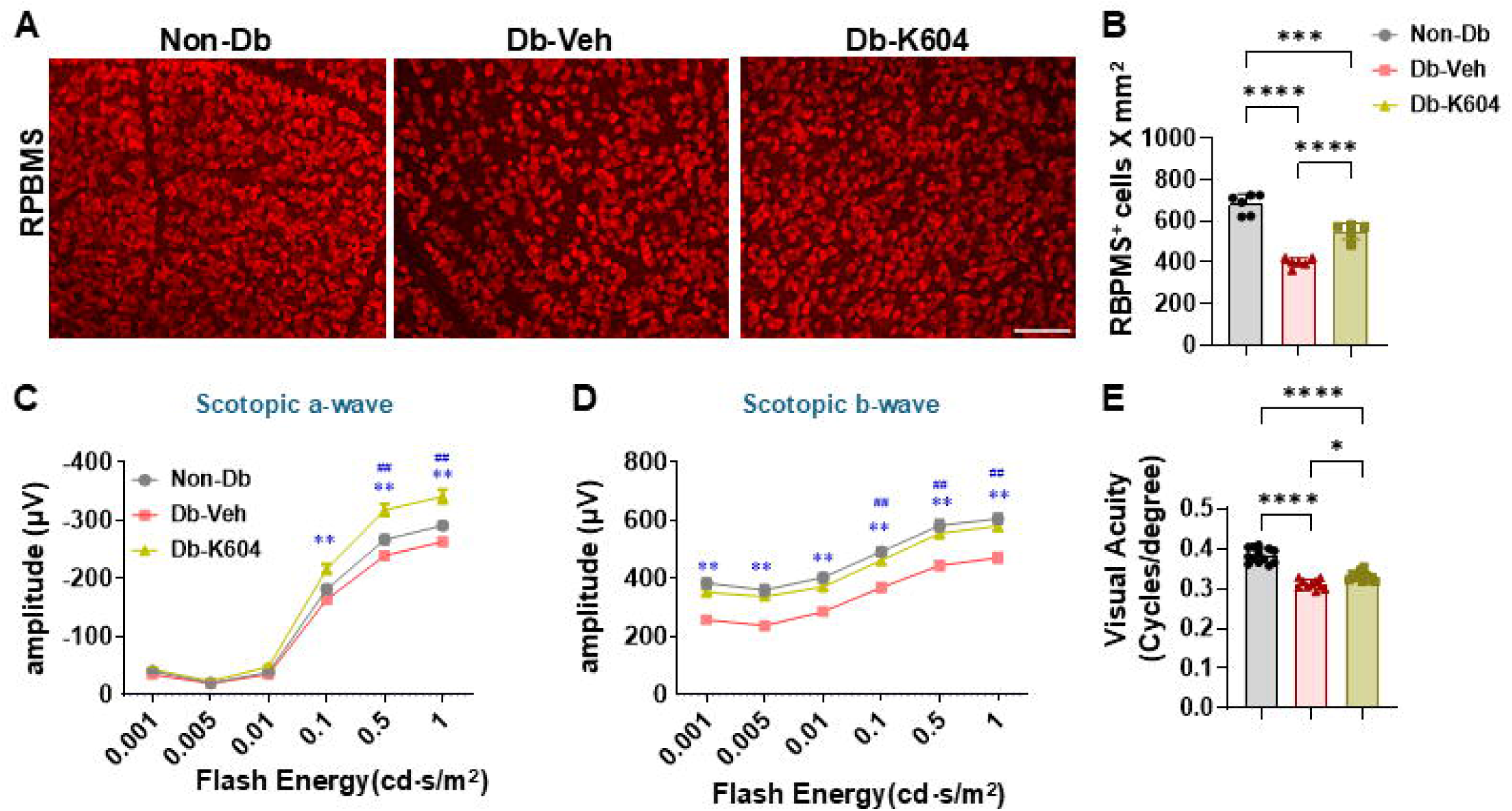
Inhibition of ACAT1/SOAT1 limits retinal ganglion cell loss and improves visual function in Ins2^Akita^ diabetic mice. Ten-week-old male diabetic Ins2^Akita^ mice were treated with ACAT1/SOAT1 inhibitor, K604 (10 mg/Kg, ip), every 2 days for 2 weeks. After 2 weeks, retinas were collected for analysis **A-B.** Confocal imaging of retinal flat mounts labeled with the neuronal marker, RBPMS and quantification showed a significant decrease in RPBMS positive retinal ganglion cells in vehicle-treated Ins2^Akita^ diabetic retinas compared to the non-diabetic control retinas. This retinal ganglion cell loss was markedly reduced by the K604 treatment. n=5-6; scale bar=40 μm. Data are presented as mean ± SEM. **C-D**. Retinal function was measured by electroretinography (ERG). Both scotopic a-wave and scotopic b-wave were improved in K604-treated Ins2^Akita^ diabetic mice as compared with the vehicle treated diabetic mice. n=12–26. Data are presented as mean ± SEM. **p<0.01 non-diabetic versus vehicle-treated diabetic mice; p<0.01 vehicle-treated diabetic versus K604-treated diabetic mice. **E**. Measurement of visual acuity by optoMotry responses (OMR) showed a marked improvement in the K604-treated diabetic mice as compared with the vehicle-treated diabetic mice. n=10-13. Data are presented as mean ± SEM. *p<0.05; ***p<0.0001.

### ACAT1/SOAT1 inhibition preserves retinal microvascular integrity in early diabetes

Diabetic mice exhibited characteristic microvascular abnormalities, including increased leukostasis, vascular permeability, and acellular capillary formation, all of which were prevented by ACAT1/SOAT1 inhibition. Quantitative analysis of attached leukocytes within the retinal vessels showed enhanced leukocyte adhesion in diabetic mice and this increase was normalized by K604 treatment (**Fig. 6A, B**). Vascular leakage, assayed by FITC-BSA fluorescence, was markedly elevated in diabetic retinas but was strongly suppressed following inhibitor treatment (**Fig. 6C, D**). Furthermore, K604 significantly blocked the diabetes-induced increase in acellular capillaries, a hallmark of diabetic microvascular degeneration (**Fig. 6E, F**). These findings demonstrate that ACAT1/SOAT1 activity contributes to retinal vascular pathology in diabetes and that its inhibition preserves microvascular integrity.

**Fig. 6.**
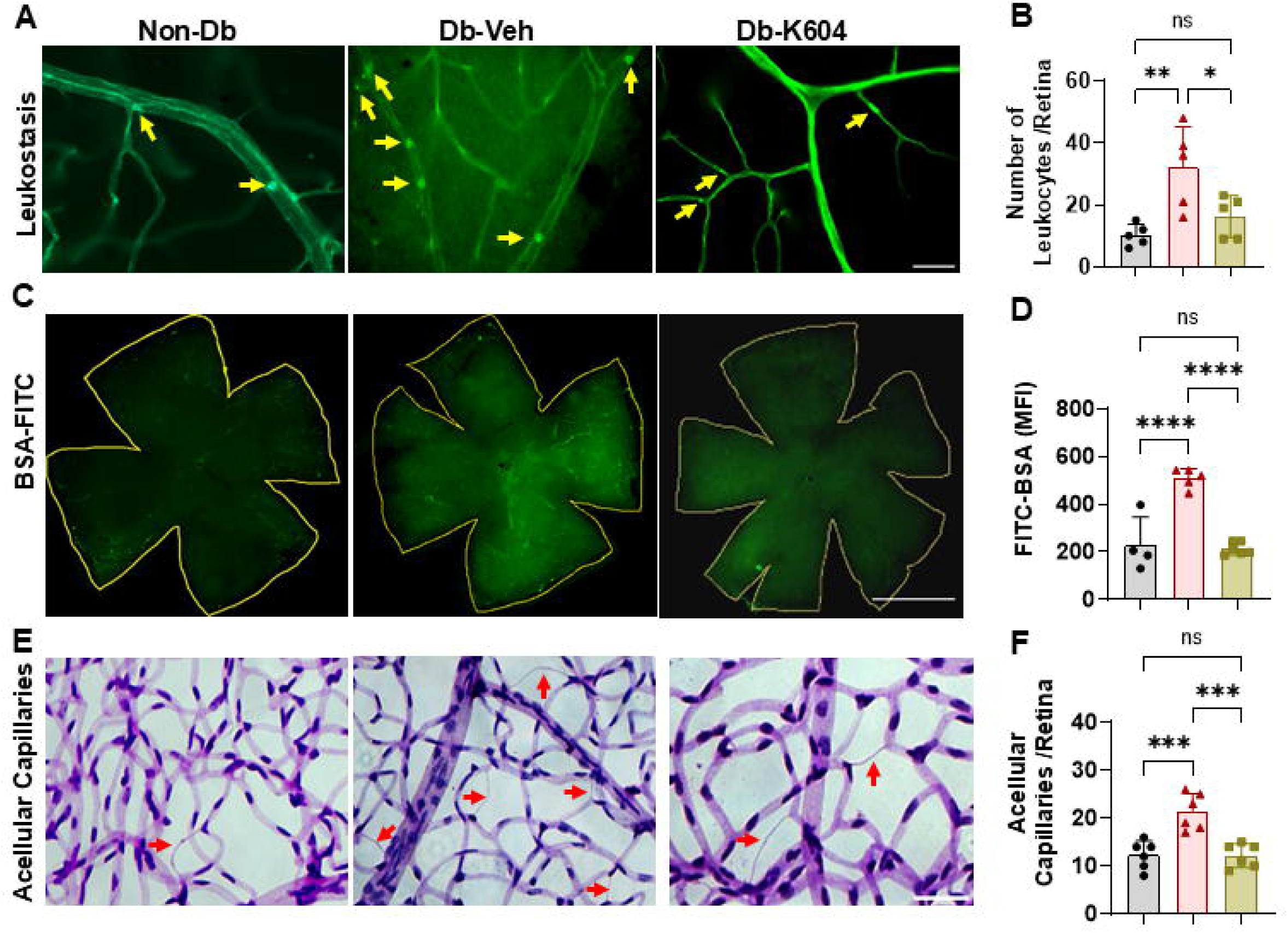
Inhibition of ACAT1/SOAT1 prevents diabetes-induced microvascular abnormalities (leukostasis, vascular permeability and acellular capillaries) in Ins2^Akita^ diabetic mice. Ten week old male diabetic Ins2^Akita^ mice were treated with ACAT1/SOAT1 inhibitor, K604 (10 mg/Kg, ip), every 2 days for 2 weeks. After the 2 weeks treatment, retinas were collected for analysis. **A-B.** Quantitative analysis shows diabetes-induced increases in leukocyte adhesion in Ins2^Akita^ retinal vessels, which were significantly prevented by K604 treatment as compared with vehicles. Data are presented as mean ± SEM. n=5-6; *p<0.05; **p<0.01; Scale bar=50 μm. **C-D**. Vascular leakage was visualized and quantified by measuring the fluorescence intensity of FITC-conjugated BSA. K604 treatment significantly prevented vascular leakage in Ins2^Akita^ mice as compared with vehicle controls. Data are presented as mean ± SEM. n=5-6; scale bar=300 μm; ****p<0.0001. **E**. K604 treatment significantly prevented diabetes-induced increases in acellular capillary formation as compared with the vehicle controls. Data are presented as mean ± SEM. n=6, scale bar=40 μm; ****p<0.001.

### ACAT1/SOAT1 inhibition limits vascular leakage and preserves visual function in later stage diabetic retinopathy

To determine whether ACAT1/SOAT1 inhibition is effective at later stages of disease, 8-month-old Ins2^Akita^ mice were treated with K604 for 2 months (**Supplementary Fig. 4A**). As expected, diabetes caused a reduction in body weight, which was not altered by K604 treatment (**Supplementary Fig. 4B**). Blood glucose levels significantly increased in diabetic mice compared with WT controls and remained unchanged following K604 treatment (**Supplementary Fig. 4C**). Oil Red O (**Fig. 7A, B**) and filipin staining (**Fig. 7C, D**) revealed pronounced lipid accumulation and cholesterol ester deposition in the ganglion cell layer, horizontal cell layers and photoreceptor outer segments of diabetic retinas. These findings were associated with high levels of 4HNE in the same location as CE deposition, indicating increases in lipid peroxidation. These alterations were significantly reduced by ACAT1/SOAT1 inhibitor treatment **(Fig.7E, F).**

**Fig. 7.**
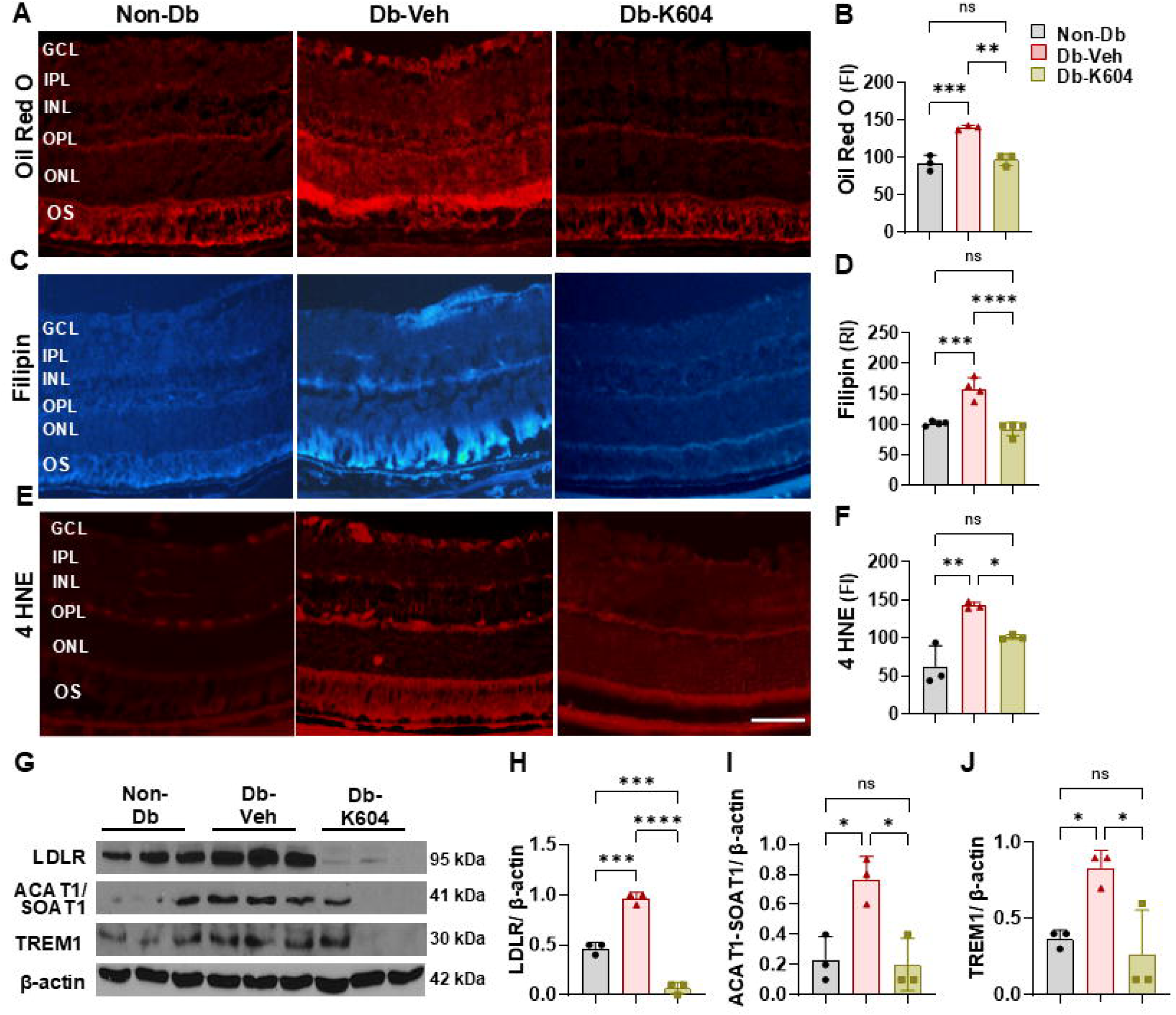
Inhibition of ACAT1/SOAT1 limits the progression of later stage DR-induced lipid/cholesterol accumulation and inflammation. Eight-months-old male Ins2^Akita^ diabetic mice were treated with ACAT1/SOAT1 inhibitor, K604 (10 mg/Kg, ip), every 2 days for 2 months. After the 2 months treatment, retinas were collected for analysis. Oil red O staining (**A, B**), filipin staining (**C, D**) and 4HNE staining (**E, F**) of frozen sections shows a strong reaction in areas of the ganglion cell layer (GCL) and photoreceptor outer segments (OS) of the DR retina, indicating lipid accumulation, CE formation, and oxidative stress, respectively. Treatment with the ACAT1/SOAT1 inhibitor, K604, significantly blocked these effects. Scale bar=25 μm. Data are presented as mean ± SEM; n=3-4; **p<0.01; ***p<0.001; ****p<0.0001. **G-J.** Western blot analysis of retinal samples from the Ins2^Akita^ diabetic mice showed marked increases in LDLR, ACAT1/SOAT1, and TREM1, as compared with the WT non-diabetic controls. These increases were normalized by treatment with K604. Mean ± SEM; n=3; *p<0.05; ***p<0.001; ****p<0.0001.

Western blot analyses further demonstrated marked increases in LDLR, ACAT1/SOAT1, and TREM1 expression in diabetic retinas compared with non-diabetic controls, all of which were normalized following K604 administration (**Fig. 7G-J)**). These results show that inhibiting ACAT1/SOAT1 reduces lipid/cholesterol accumulation and inflammatory/oxidative signaling even at advanced stages of diabetic retinopathy.

### K604 preserves vascular integrity and vision in late-stage DR

We determined whether long-term ACAT1/SOAT1 inhibition preserves retinal vascular integrity and function in advanced diabetic retinopathy. Albumin staining of retinal sections revealed extensive vascular leakage surrounding retinal vessels positive for isolectin B_4_ in diabetic retinas. This was markedly reduced by K604 treatment (**Fig. 8A, B**). Consistent with improved vascular integrity, optomotor response testing demonstrated a significant preservation of visual acuity in K604-treated diabetic mice compared with vehicle-treated controls (**Fig. 8C**). Functional assessment by ERG further showed preservation of the scotopic a- and b-wave amplitudes in inhibitor-treated mice, indicating preserved photoreceptor and bipolar cell function (**Fig. 8D, E**). These findings demonstrate that ACAT1/SOAT1 inhibition not only reduces vascular leakage but also preserves visual function in later stages of diabetic retinopathy.

**Figure 8.**
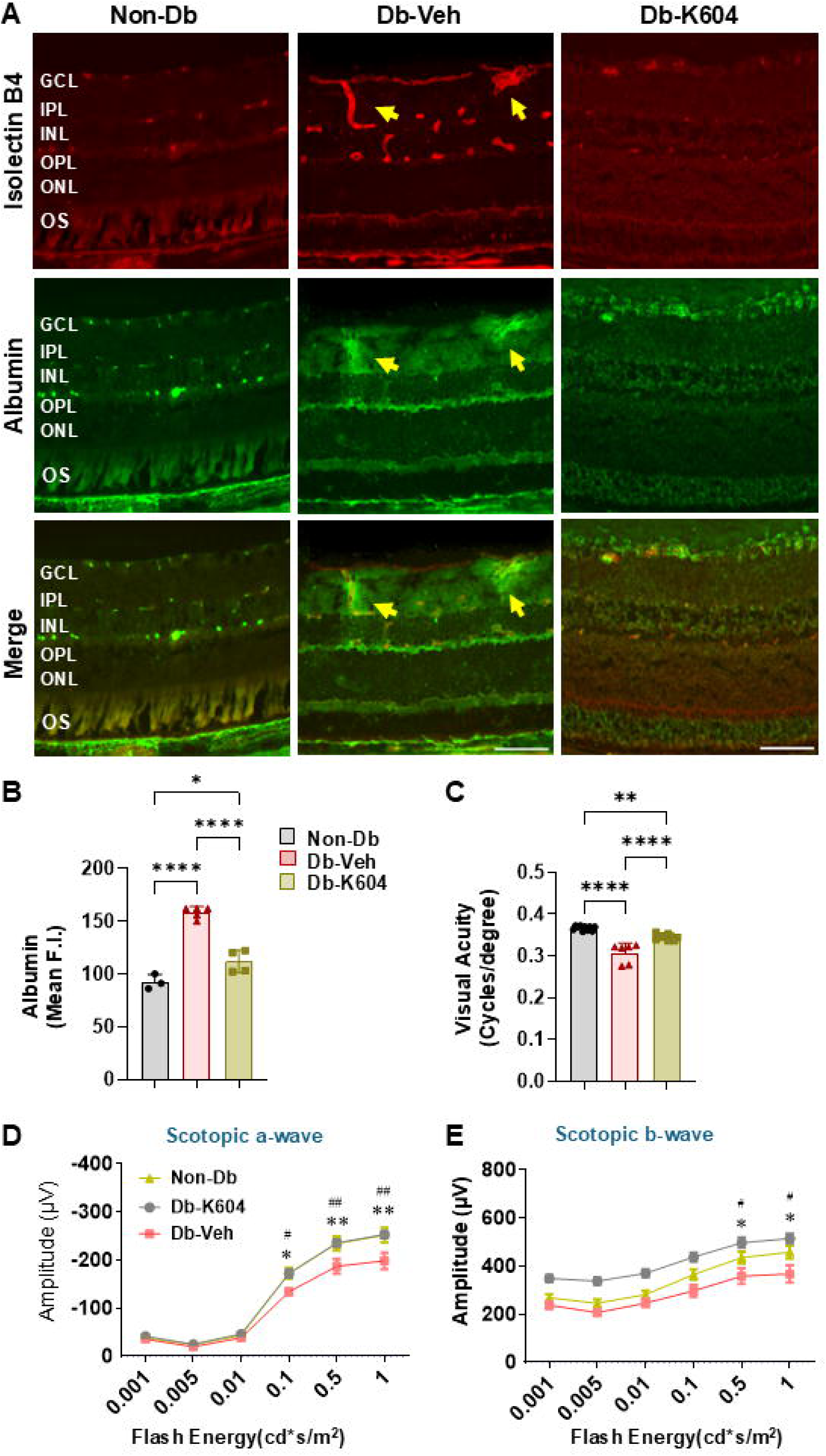
ACAT1/SOAT1 inhibitor treatment limits vascular leakage and preserves visual function in later stage DR. Eight-months old male Ins2^Akita^ diabetic mice were treated with ACAT1/SOAT1 inhibitor, K604 (10 mg/Kg, ip), every 2 days for 2 months. After 2 months, retinas were collected for analysis. **A-B**. Albumin staining of retina sections showed a strong reaction in areas of Isolectin B4-positive blood vessels in the diabetic retina, indicating vascular leakage. Treatment with ACAT1/SOAT1 inhibitor, K604, significantly blocked this effect. Scale bar=25 μm; data are presented as mean ± SEM; n=3-4; *p<0.05; ****p<0.0001. **C**. Measurement of visual acuity by optoMotry responses (OMR) showed a significant impairment in the vehicle-treated diabetic Ins2^Akita^ mice that was significantly blunted in the K604-treated diabetic mice. n=6-10. Data are presented as mean ± SEM; **p<0.01; ****p<0.0001. **D-E**. Electroretinography (ERG) showed improvement in both scotopic a-wave and scotopic b-wave in K604-treated diabetic mice as compared with the vehicle treated diabetic mice. n=6-10. Data are presented as mean ± SEM. **p<0.01 non-diabetic versus vehicle-treated diabetic mice; p<0.01 vehicle-treated diabetic versus K604-treated diabetic mice.

## Discussion

Our studies demonstrate that diabetes induces increases in CE formation due to activation of ACAT1/SOAT1, which in turn induces key components of DR, including increases in oxidative stress, inflammation, vascular pathology, and neuronal dysfunction. Using the Ins2^Akita^ model of type 1 diabetes, we show that systemic treatment with the specific ACAT1/SOAT1 inhibitor K604 prevents CE accumulation, normalizes inflammatory and angiogenic signaling, protects against microvascular and neuronal injury, and preserves visual function. Importantly, these effects were observed in both early- and late-stage intervention paradigms, highlighting the therapeutic potential of targeting cholesterol esterification across the spectrum of DR duration and progression.

Our findings provide mechanistic insight into how dysregulated cholesterol metabolism contributes to diabetic retinal pathology. We observed strong upregulation of ACAT1/SOAT1 and LDLR in diabetic retinas, consistent with increased cholesterol uptake and esterification. The increase in CE formation was associated with increases in oxidative stress, glial activation, and elevated pro-inflammatory mediators including TREM1, MCSF, IL-1β, TNF-α, and VEGF. Previous studies in models of atherosclerosis have shown that dyslipidemia triggers increased TREM1 expression on myeloid cells. TREM1 is a potent amplifier of inflammatory signals (*21, 22*). It has been shown to trigger monopoiesis, production of pro-atherogenic cytokines, and foam cell formation in atherosclerotic lesions (*23*). Our findings show that increases in ACAT1/SOAT1 activity, CE formation, and expression of TREM1 are linked to oxidative stress, upregulation of inflammatory mediators, and retinal neurovascular dysfunction in the Ins2^Akita^ diabetic mice retina demonstrates a novel role of this pathway in DR.

This CE-driven inflammatory state aligns with previous reports linking dyslipidemia to DR severity and provides a mechanistic explanation for the limited efficacy of systemic lipid-lowering therapies in clinical trials. Unlike systemic statins or fibrate targeting therapies, ACAT1/SOAT1 inhibition directly targets intracellular cholesterol esterification within the retina, thereby reducing the pool of bioactive lipids that fuel oxidative stress and inflammation. Identification of the specific mechanisms by which increased CE formation promotes oxidative stress and expression of TREM1, and other inflammatory mediators require further investigation. Studies in models of cardiovascular disease have shown that oxidized CE can induce the activation of several inflammatory mediators (*24*). While this mechanism is clearly established in cardiovascular disease, its role in DR is completely unknown.

We also establish a critical link between cholesterol esterification and vascular dysfunction. K604 treatment prevented diabetes-induced leukostasis, vascular leakage, and acellular capillary formation, suggesting that CE accumulation contributes to endothelial injury and blood–retina barrier breakdown. These results are consistent with prior work implicating lipid peroxidation and inflammatory signaling in retinal vascular dysfunction. Moreover, by reducing VEGF expression, ACAT1/SOAT1 inhibition addresses one of the key drivers of pathological neovascularization in proliferative DR.

Beyond vascular effects, we demonstrate that ACAT1/SOAT1 inhibition provides robust neuroprotection. K604 treatment preserved retinal ganglion cell survival, improved ERG responses, and enhanced visual acuity. This is particularly significant, as visual dysfunction often precedes clinically detectable vascular lesions in DR. By targeting upstream metabolic dysregulation, ACAT1/SOAT1 inhibition may intervene earlier in the disease course than current anti-VEGF therapies, which primarily address late stage neovascular complications.

The protective effects of K604 were observed without changes in either systemic glucose levels or body weight, confirming that the benefits were independent of glycemic control. We were not expecting K604 to normalize glucose levels since Ins2^Akit^ mice undergo a spontaneous mutation of the *Ins2* gene resulting in early onset diabetes due to lack of insulin production (*25*). However, further studies are needed to assess the effect of K604 on plasma glucose levels in different models of diabetes. This is relevant clinically, as intensive glycemic control is often insufficient to fully prevent progression of DR. Our findings suggest that adjunctive therapy targeting cholesterol esterification complementing current glucose-lowering and anti-VEGF treatments may be helpful in limiting DR.

There are several potential limitations to consider. First, our study was performed exclusively in Ins2^Akita^ mouse, a model of type 1 diabetes, and it will be important to confirm these findings in models of type 2 diabetes and in human retinal tissue. Second, while K604 is a selective ACAT1/SOAT1 inhibitor, off-target effects cannot be fully excluded and development of more retina-specific ACAT1/SOAT1 inhibitors would be desirable for clinical translation. Finally, long-term safety studies are needed to determine whether chronic ACAT1/SOAT1 inhibition alters other cholesterol-dependent pathways in the retina or peripheral tissues.

In conclusion, our work identifies ACAT1/SOAT1 as a key regulator of lipid-driven pathology in diabetic retinopathy and establishes cholesterol esterification as a novel therapeutic target. By reducing oxidative stress, inflammation, vascular and neuronal dysfunction, ACAT1/SOAT1 inhibition offers a multifaceted approach to DR treatment. These findings provide a strong rationale for further preclinical development of ACAT1/SOAT1 inhibitors and may open new avenues for treating both early and advanced stages of diabetic retinopathy.

### Significance Statement

Diabetic retinopathy (DR) remains the leading cause of blindness in working-age adults, and current therapies, including anti-VEGF injections, primarily target late-stage vascular complications without addressing the underlying metabolic drivers of disease. Our study identifies cholesterol esterification via ACAT1/SOAT1 as a central metabolic mechanism linking diabetes to oxidative stress, inflammation, vascular injury, and neuronal dysfunction in the retina. Pharmacologic inhibition of ACAT1/SOAT1 not only reduced vascular leakage and inflammation but also preserved visual function, even when treatment was initiated at advanced disease stages. These findings highlight cholesterol esterification as a novel, targetable pathway in DR pathogenesis and provide preclinical proof-of-concept for ACAT1/SOAT1 inhibitors as adjunctive or alternative therapies to current treatment strategies.

## METHODS

### Human donor retinas

Retinal tissue sections from human donors with proliferative diabetic retinopathy (PDR) and age matched non-diabetic controls were purchased from National Disease Research Interchange (NDRI, Philadelphia, PA). Sections were deparaffinized and immunolabeled as described in the immunolocalization section. Patient information is provided in **supplementary table 1**.

### Human vitreous samples

Samples were collected from patients undergoing pars plana vitrectomy (PPV) due to proliferative diabetic retinopathy (PDR) with retinal detachment and patients undergoing PPV for reasons other than PDR (chronic retinal detachment, open globe injury, or macular hole). Vitrectomy samples were concentrated using Amicon Ultra 30k centrifuge filter devices (cat # UFC903008) and CE levels were measured as previously described (*26*). Patient information is provided in **supplementary table 2**.

### Mouse model of type 1 diabetes and drug treatment

Experiments were performed in accordance with the ARVO Statement for the Use of Animals in Ophthalmic and Vision Research and were approved by the institutional animal care and use committee animal welfare. Ins2Akita Type 1 diabetic mice and littermate control mice were bred in our facility. Mice were sacrificed at different ages for experimental endpoints and littermates from the same cage were compared. Diabetes was confirmed by blood glucose measurement and genotyping as previously described (*27*),(*28*). Mice were treated with intraperitoneal injections of a specific ACAT1/SOAT1 inhibitor (K604, MedChem Express, 10 mg/Kg in PBS) or vehicle control (PBS) every 2 days. One group of mice was treated for 2 weeks beginning at 10 weeks of age and sacrificed at 3 months. Another group of mice was treated for 2 months beginning at 8 months and sacrificed at 10 months. Previous studies have shown that intraperitoneal injection of K604 is safe and that the drug can penetrate the blood-retinal barrier (*18*).

### Western blotting

Lysates from retina samples and cell lines were prepared for protein quantification and western blot analysis following our established protocol (*29*). The samples were homogenized in modified RIPA buffer (20 mM Tris–HCl, 2.5 mM EDTA, 50 mM NaF, 10 mM Na4P2O7, 1% Triton X-100, 0.1% sodium dodecyl sulfate, 1% sodium deoxycholate, 1 mM phenylmethylsulfonylfluoride, pH 7.4). Samples containing equal amounts of protein were separated by 10% or 12% sodium dodecyl sulfate polyacrylamide gel electrophoresis, transferred to polyvinylidene difluoride (PVDF) or nitrocellulose membrane, and reacted overnight with anti-LDLR, anti-ACAT1/SOAT1, anti-TREM1, anti-MCSF1, anti-TNF-a, and anti-VEGF antibodies in 5% milk or 2% BSA, followed by incubation with corresponding horseradish peroxidase-linked secondary antibodies. Bands were quantified by densitometry, and the data were analyzed using ImageJ software. Equal loading was verified by stripping the membranes and reproving them with a mouse monoclonal antibody against β actin. Antibodies were obtained from various sources. Their catalog information, together with working dilutions, is provided in **supplementary table 3**.

### Quantitative real time RT PCR (qRT PCR) analysis

Total RNA from mouse retina samples was extracted with an RNAqueous 4PCR total RNA isolation kit (Invitrogen, Carlsbad, CA, US), and qRT PCR was performed as described previously (*18, 30*). Briefly, a 0.25 μg sample of RNA was utilized as a template for reverse transcription using M-MLV reverse transcriptase (Invitrogen). qRT PCR was performed on an ABI 7500 Real Time PCR System (Applied Biosystems, Foster City, CA) with the respective gene-specific primers listed in **supplementary table 4**. The relative mRNA expression was calculated using the comparative threshold cycle (ΔΔCt) method against the internal control, hypoxanthine phosphoribosyl-transferase (HPRT). Expression levels for all genes are reported as fold change to room air controls.

### Immunolocalization

Retinal frozen sections and flat mounts were processed for immunolabeling according to our standard protocol (*18, 31*). The samples were blocked with normal goat serum or donkey serum and incubated overnight with IB**_4_**, anti-Iba1, anti-GFAP, anti-ACAT1/SOAT1, and anti-4HNE antibodies. The samples were washed 3 times with PBS and incubated with secondary antibodies (Invitrogen, Waltham, MA). Antibodies were obtained from various sources, and their catalog information, together with working dilutions, is provided in supplementary table 2. Tissue sections were also reacted with secondary antibody alone to rule out non-specific reactivity. Images were captured using a Zeiss Axioplan2 fluorescence microscope or Zeiss 780 inverted Confocal microscope (Carl Zeiss Meditec, Inc. Dublin, CA).

### Oil Red O staining for detection of neutral lipids

Cholesterol and other neutral lipids were detected as described previously (*18, 32*). Retina frozen sections were reacted with 0.5% Oil Red O (Sigma-Aldrich, St. Louis, MO, dissolved in 1,2-isopropanol, 15 min, room temperature), rinsed in 60% 2-propanol for 5 min and rinsed in distilled water (twice, 5 min each time). Sections were mounted in aqueous mounting media to capture the images using a Zeiss Axioplan 2 fluorescence microscope (Carl Zeiss Meditec, Inc., Dublin, CA) equipped with a 420 nm excitation filter, 520 nm barrier filter, and a 20× lens.

### Filipin staining for detection cholesterol ester

Retinal frozen sections were reacted with the fluorescent polyene antibiotic filipin (Sigma-Aldrich, St. Louis, MO) to detect CE. This compound binds specifically to sterols and interacts with the 3-β hydroxyl group of cholesterol. For CE detection, we followed the protocol described by Rudolf and Curcio (*33*). Native unesterified cholesterol (UC) was extracted from cryosections by rinsing in 60% ethanol for 10 min. Native CE was hydrolyzed with cholesterol esterase (1.65 units/mL) in PBS for 3 h at 37 °C (C9281, Sigma St. Louis, MO). The newly released UC by the hydrolysis of CE was stained with 50 ng/mL filipin in PBS for about 60 min in the dark at room temperature.

### Cholesterol ester measurement by ELISA

Levels of CE formation were determined as described previously (*18*) using the luminescence-based Cholesterol/Cholesterol Ester Glo™ Assay (J3190, Promega, Madison, WI) according to the manufacturers’ protocol. Briefly, human vitreous samples and mouse retinas were collected and homogenized, and blood plasma was collected. All samples were diluted (1:10) in cholesterol lysis solution and incubated at 37 °C for 30 min. Equal amounts of extracts were incubated in cholesterol detection reagent with and without cholesterol esterase enzyme in 96 well white bottom plates at room temperature for 1 h. Luminescence was recorded using a Polar Star Omega microplate reader (BMG Labtech Inc, NC). Total and free cholesterol concentrations were measured by comparing the luminescence of samples with and without cholesterol esterase, respectively. Concentrations of CE were calculated as the difference between total and free cholesterol concentrations.

### ROS production

The dihydroethedium (DHE) method was used to evaluate superoxide formation as described previously (*34, 35*). Briefly, fresh frozen sections were preincubated with or without SOD-polyethylene glycol (400 U/ml, Sigma-Aldrich, St. Louis, MO, USA) for 30 min, followed by reaction with DHE (10 μM) for 15 min at 37 °C. Superoxide, the absence of SOD, oxidizes DHE to ethidium bromide, which binds to DNA and fluoresces red (*36*). A fluorescence microscope (AxioVision; Carl Zeiss) was used to obtain the DHE images immediately after incubation. DHE was excited at 488 nm with an emission spectrum of 610 nm. Six images per slide were taken and the relative fluorescence intensity was measured by ImageJ software.

### Isolation of the retinal vasculature and measurement of acellular capillaries

The trypsin digestion method was used for isolating the retinal vasculature as previously described with minor modification(*35*). Eyeballs were removed and fixed with 4% paraformaldehyde overnight. Retinas were carefully dissected and incubated in distilled water at room temperature with gentle shaking for at least 24 h. Then the retinas were digested with 3% trypsin (Difco Trypsin 250, Becton Dickinson and Company, Sparks, MD, USA) in 0.1 M Tris buffer (pH 7.8) for 1.5 h at 37°C on an orbital shaker (50 r.p.m.). Retinas were washed carefully in several changes of fresh distilled water until no non-vascular tissue was observed under the microscope. The isolated retinal vasculature was air-dried on silane-coated slides and stained with periodic acid–Schiff and hematoxylin. Acellular capillaries were counted in 10 random areas of the mid-retina. The field area was calculated using ImageJ software. The numbers of acellular capillaries were divided by the field area to get number of acellular capillaries per/mm^2^ of retina.

### Retinal vascular permeability

Retinal vascular permeability was tested by evaluating albumin extravasation into the retinal parenchyma using albumin staining on retinal cross sections as described previously (*29*). Mice received intraperitoneal injections of 10 mg/Kg bovine serum albumin (BSA)–Alexa-Fluor 488 conjugate (Invitrogen-Molecular Probes, Eugene, OR). After 2 hours, a thoracotomy was performed on deeply anesthetized mice. The right atrium was cut, and a heparinized catheter was inserted into the left ventricle. Mice were perfused transcardially with PBS 1X at 37° C for 2 minutes to wash out the BSA-Alexa-Fluor 488 from the retinal vessels. Eyeballs were enucleated and fixed overnight in 4% paraformaldehyde in PBS at 4 °C. The retinas were dissected into whole mounts and images were acquired using a fluorescence microscope (Keyence, BZ-X700). The images of the whole retinal flat mounts were constructed by capturing a series of 6 pictures from all samples using a 4× lens. Fluorescence intensity was quantified using Image J.

Other eyeballs were embedded in OCT compound for preparation of frozen sections which were reacted with a rabbit anti-mouse albumin antibody (1:400, Bethyl Laboratory) followed by reaction with Oregon green labeled goat anti-rabbit antibody (1:500; Invitrogen). Fluorescence images were acquired using a fluorescence microscope and relative intensity of extravascular albumin was measured by ImageJ.

### Leukocyte Adhesion

Adhesion of leukocytes to the wall of the retinal vessels was evaluated as described previously(*31*). After the induction of deep anesthesia, the chest cavity was carefully opened, and a 20-gauge perfusion cannula was introduced into the aorta. Drainage was achieved by opening the right atrium. The animals were then perfused with 10 mL of phosphate-buffered saline (PBS) to wash out nonadherent blood cells. Next, the animals were perfused with 10 mL fluorescent isothiocyanate (FITC)–labeled concanavalin A (Con A) lectin (40 μg/mL in PBS, pH 7.4; Vector Laboratories, Burlingame, CA) to label the adherent leukocytes and vascular endothelial cells. Residual unbound Con A was removed by right ventricle perfusion with PBS at a rate of 1.8 ml per minute to temperature of 37^0^C. Eyeballs were enucleated and fixed overnight in 4% paraformaldehyde in PBS at 4 °C. The retinas were dissected into flat mounts, analyzed by fluorescence Keyence microscopy, and the total number of adherent leukocytes per retina was calculated

### Wholemount retina and GCL neuronal cell counting

Neuronal degeneration was assessed as previously described (*37, 38*) Eyeballs were enucleated and fixed overnight in 4% paraformaldehyde in PBS at 4 °C. Retinas were dissected into flat mounts and incubated with the neuronal marker anti-RBPMS antibody overnight at 4 °C and then incubated with Alexa fluor 488 conjugated secondary antibodies. Four images were taken in the midperiphery region of each retina flat mount using a Keyence fluorescence microscope and numbers of RBPMS-positive cells were calculated using ImageJ software. Data are represented as numbers of RBPMS-positive cells in the GCL per mm^2^.

### OptoMotry

OptoMotry (Cerebral Mechanics Inc.) was employed for screening visual function using the optokinetic tracking (OKT) response (*39, 40*) (*41*). Individual unrestrained mice were placed on an elevated platform surrounded by four computer monitors. The monitors project a virtual stimulus in the form of a sine wave that rotates around the animal. Animals track the grating with reflexive head and neck movements in the direction of grid rotation. By reversing the direction of the grid rotation, the system measures visual function in each eye separately because motion in the temporal to nasal direction of either eye elicits a tracking response. A camera monitors the behavior of the animal from above, allowing the observer to detect the mouse tracking responses in real time and give a score of yes or no. The whole study was conducted by an observer who was blinded to the treatment groups. To examine visual acuity, spatial frequency thresholds (cycles per degree) were measured by systematically increasing the spatial frequency of the grating (decreasing the bar width) at full contrast until mice no longer tracked. Rotation speed was fixed at 12 degrees per second, and the data was managed and generated by the software. Data are presented as responses from both eyes.

### Electroretinography (ERG)

ERG was performed to investigate neuronal functions using the Celeris-Diagnosys system (Diagnosys, Lowell, MA, USA) as described previously (*41*). After overnight dark adaptation, mice were anesthetized and pupils were dilated with topical 0.5% tropicamide (Akorn, Lake Forest, IL, USA) and 2.5% phenylephrine HCL (Paragon BioTeck, Portland, OR, USA). Mouse corneas were then moisturized with a thin layer of GenTeal Lubricant Eye Gel (Alcon, Ft. Worth, TX). The light guide electrodes were placed on the gel and recording began after the scotopic and photopic testing was set up. For each animal, the light guide electrode was used to present light flashes over a range of intensities and record ERG responses. The a-wave amplitude was measured as the difference between the pre-stimulus baseline and the trough of the a-wave. The b-wave amplitude was measured from the trough of the a-wave to the peak of the b-wave. The results are presented as average values of amplitude with the two eyes of each mouse.

### Statistical analysis

All data were analyzed by investigators blinded to the group allocation. Differences among the groups were compared by an independent two sample t test for two groups or one-way ANOVA followed by a post hoc test for multiple comparisons. Values are represented as means ± standard error of the means (SEM). P values of less than 0.05 were considered significant.

## List of Supplementary Materials

Supplementary figures 1 to 4 for multiple supplementary figures.

Supplementary table 1 to 4 for multiple supplementary tables.

## Acknowledgements

Dr. Ta-Yuan Chang, Ph.D. and Dr. Catherine Chang from Dartmouth Geisel School of Medicine, Hanover, New Hampshire, Dr. Eric J. Belin de Chantemèle from Vascular Biology Center Augusta University, Augusta Georgia, Dhruvi Paladiya, Medical College of Georgia, Augusta GA.

## Funding

This study was supported by grants from the National Institute of Health [R01EY035683, R01EY011766, R01EY030500, P30EY031631] and the Culver Vision Discovery Institute at Augusta University.

## Author contribution

Conceptualization: MAR

Methodology: MAR, SAHZ, RBC

Investigation: MAR, SAHZ, RBC, TL, XZ

Visualization: MAR

Funding acquisition: MAR, RBC

Project administration: RBC Supervision: MAR

Writing – original draft: SAHZ, MAR, RBC, MY

Writing – review & editing: MAR, RBC, WRC, SB, MY

## Competing interests

The authors declare that they have no competing interests.

## Data and materials availability

The data generated and/or analyzed during the current study are available from the corresponding authors on reasonable request.

## Declarations

### Ethics approval and consent to participate

All experiments were conducted in accordance with the ARVO Statement for the Use of Animals in Ophthalmic and Vision Research and were approved by the Augusta University Institutional Animal Care and Use Committee (animal protocol # 2008-0243).

Human tissues were from donors who had given consent for the use of the tissues for research purposes.

## Notes

### Competing Interest Statement

The authors have declared no competing interest.

### Summary of Updates

Updated ways to present supplementary Table 1 and supplementary Table 2. Inserting the legend of figure 8, which was not included in the previous version

