## Supplementary figures and images for "Novel Role of AcylCoA:cholesterol acyltransferase 1 (ACAT1/SOAT1) in Diabetic Retinopathy"

### Supplementary Figure 1

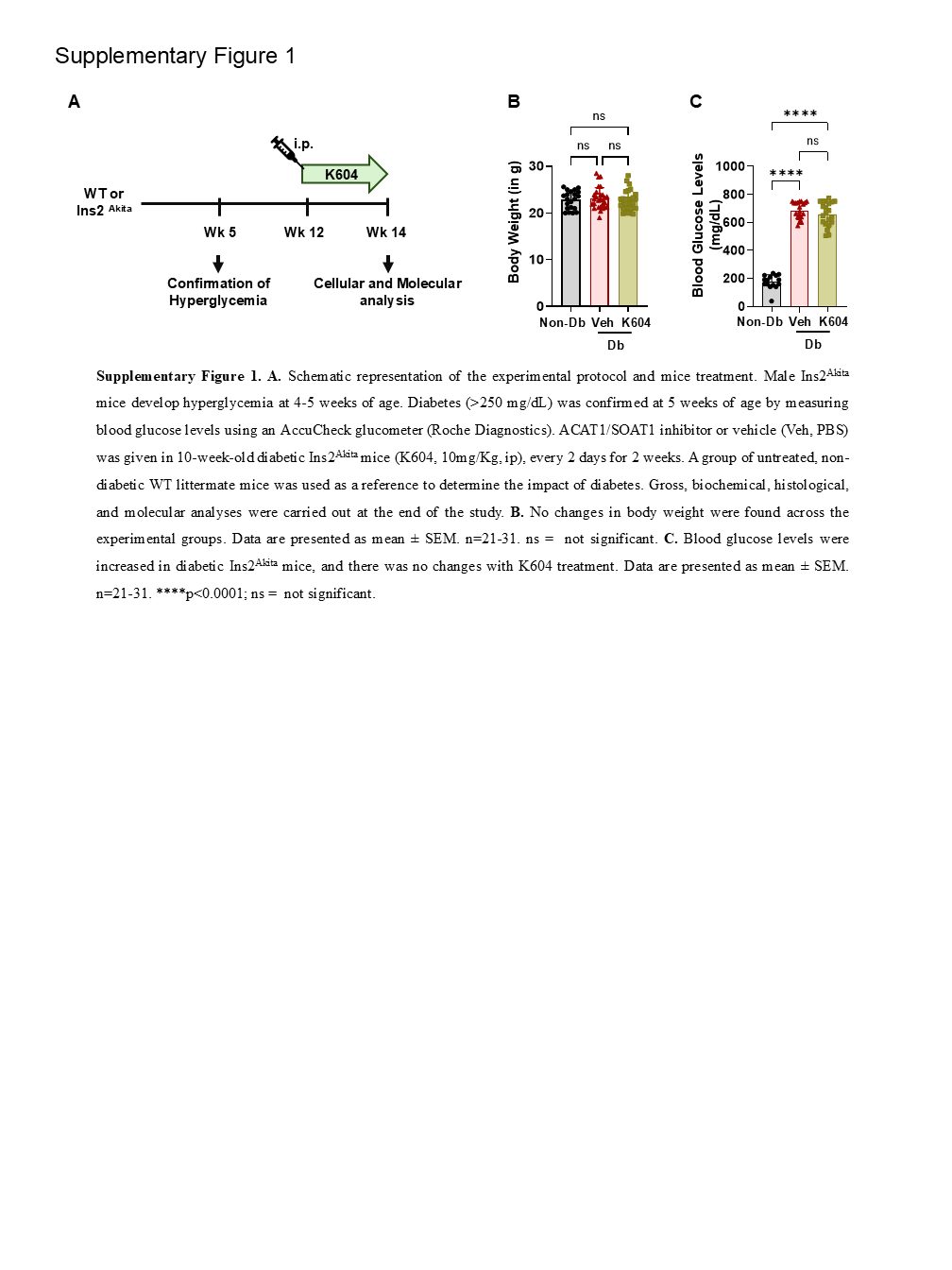

### Supplementary Figure 2

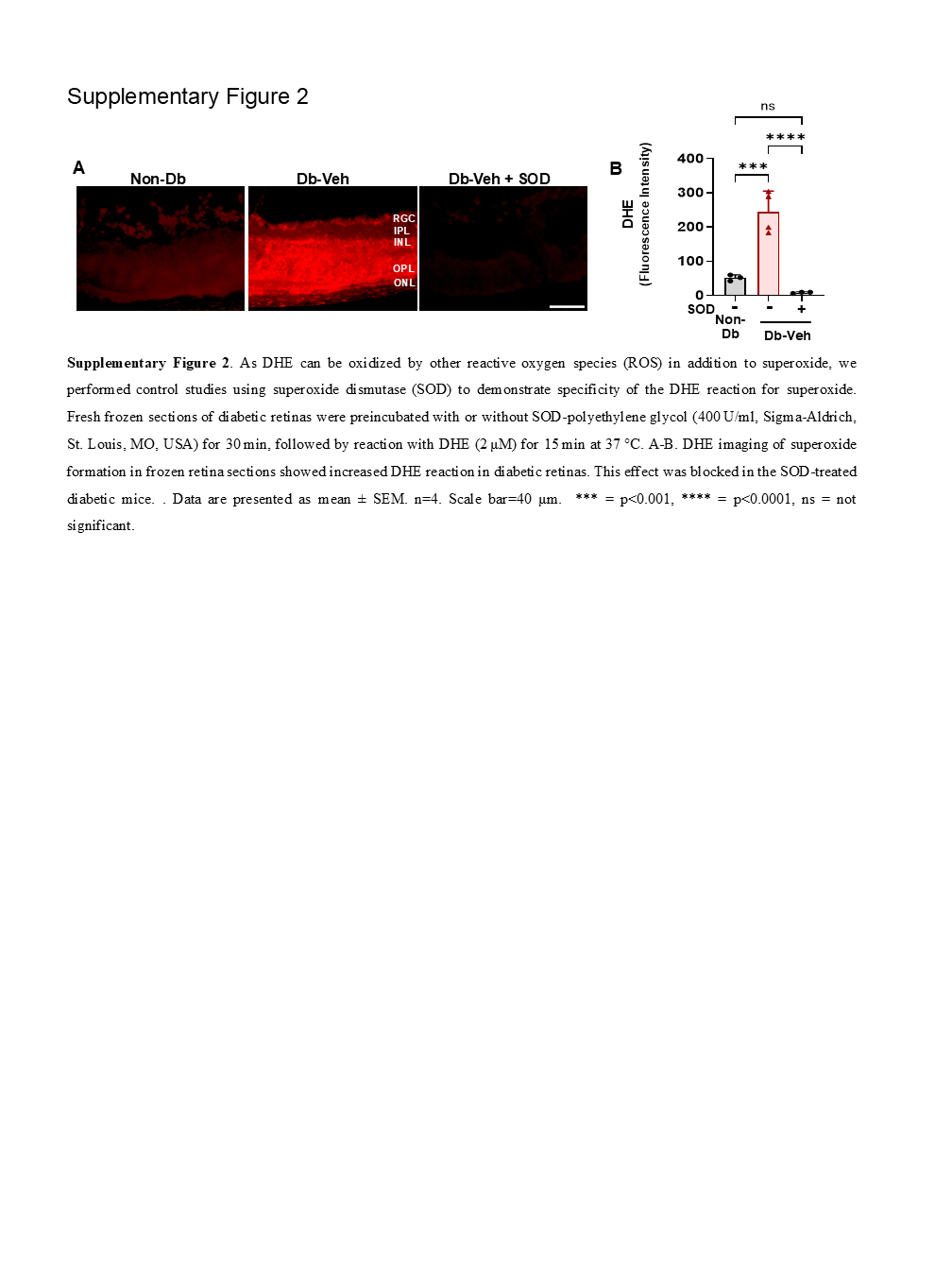

### Supplementary Figure 3

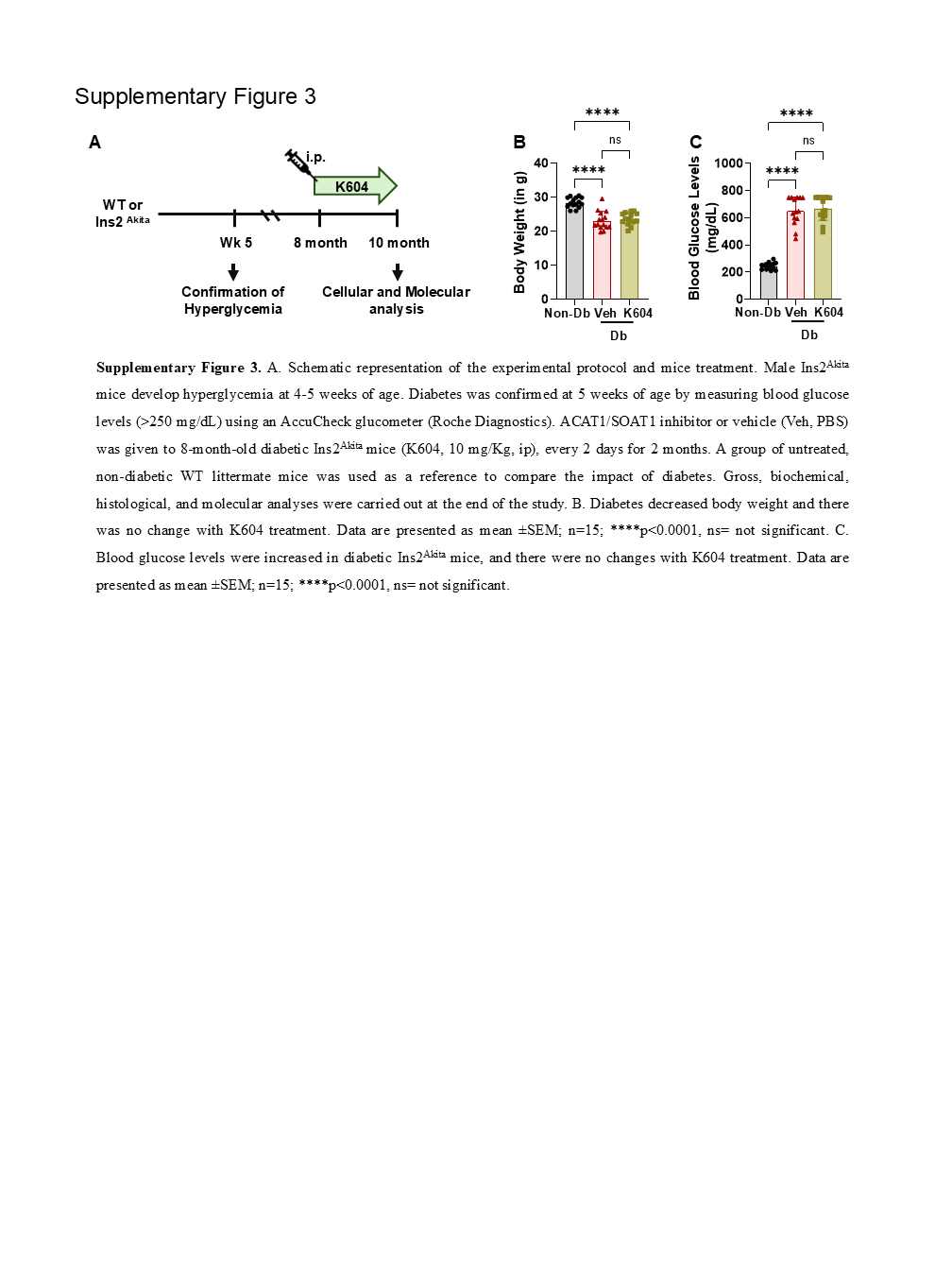

### Supplementary Figure 4

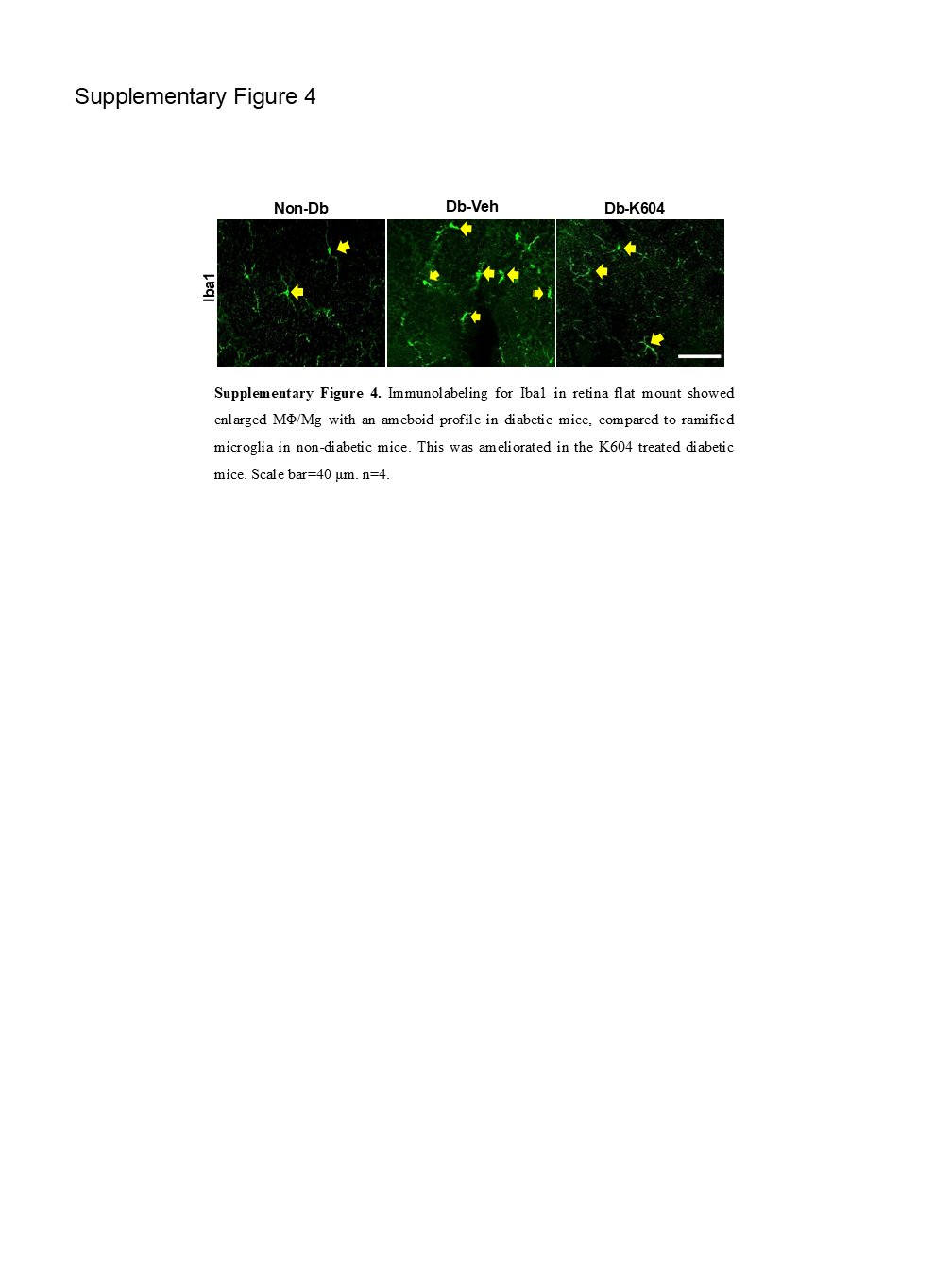

### Supplementary Table 1

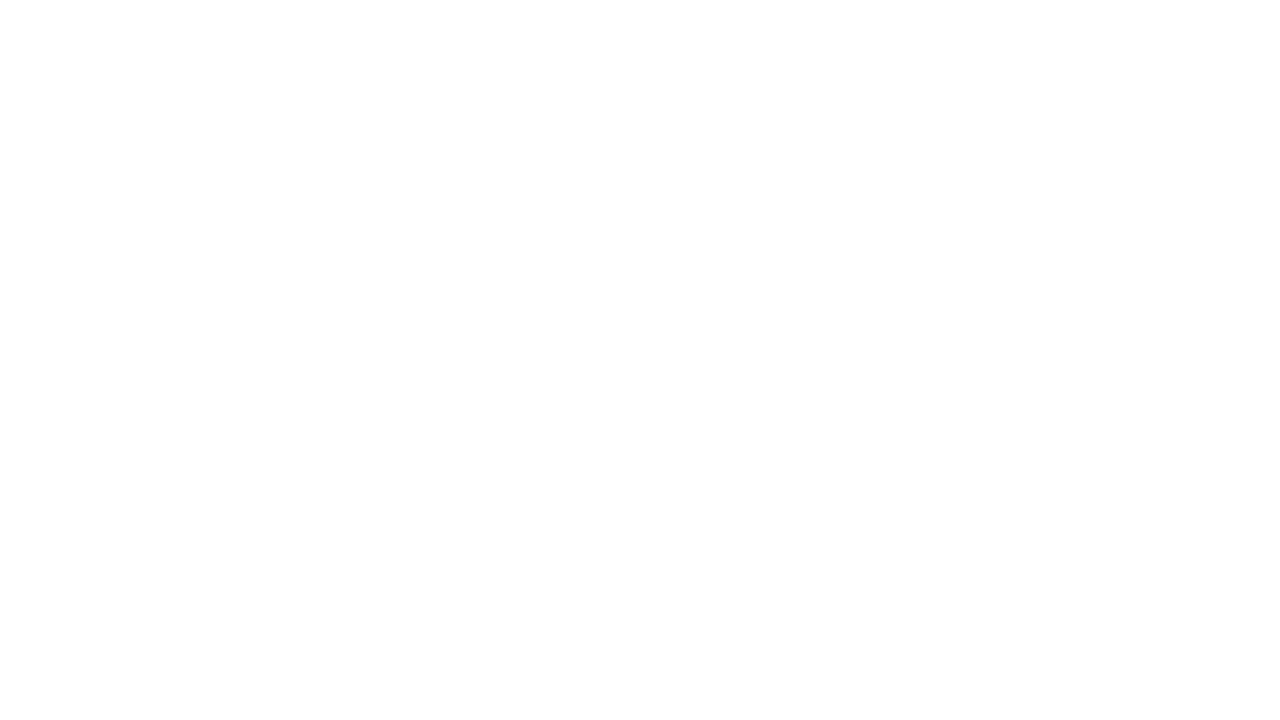

### Supplementary Table 2

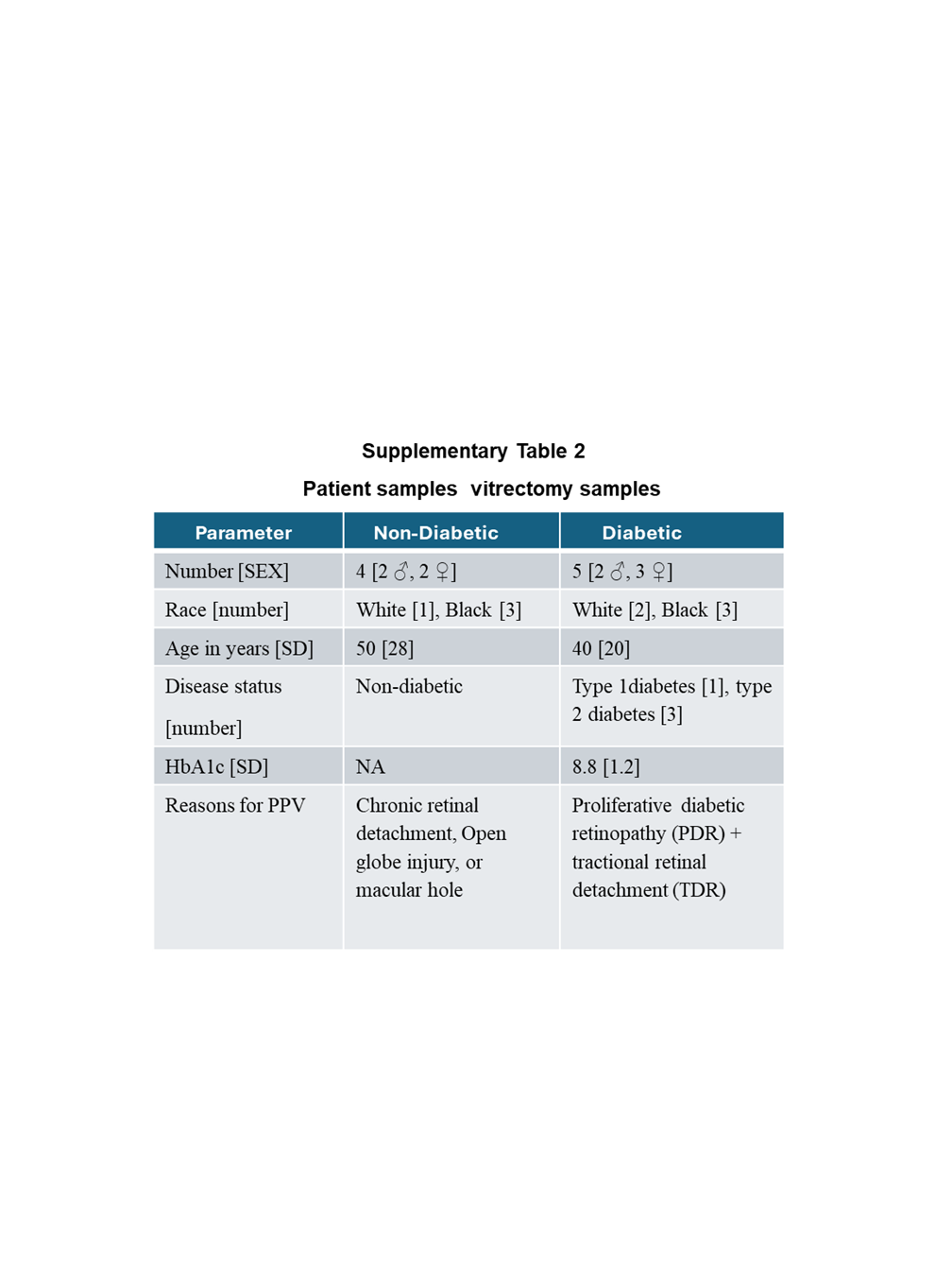

### Supplementary Table 3

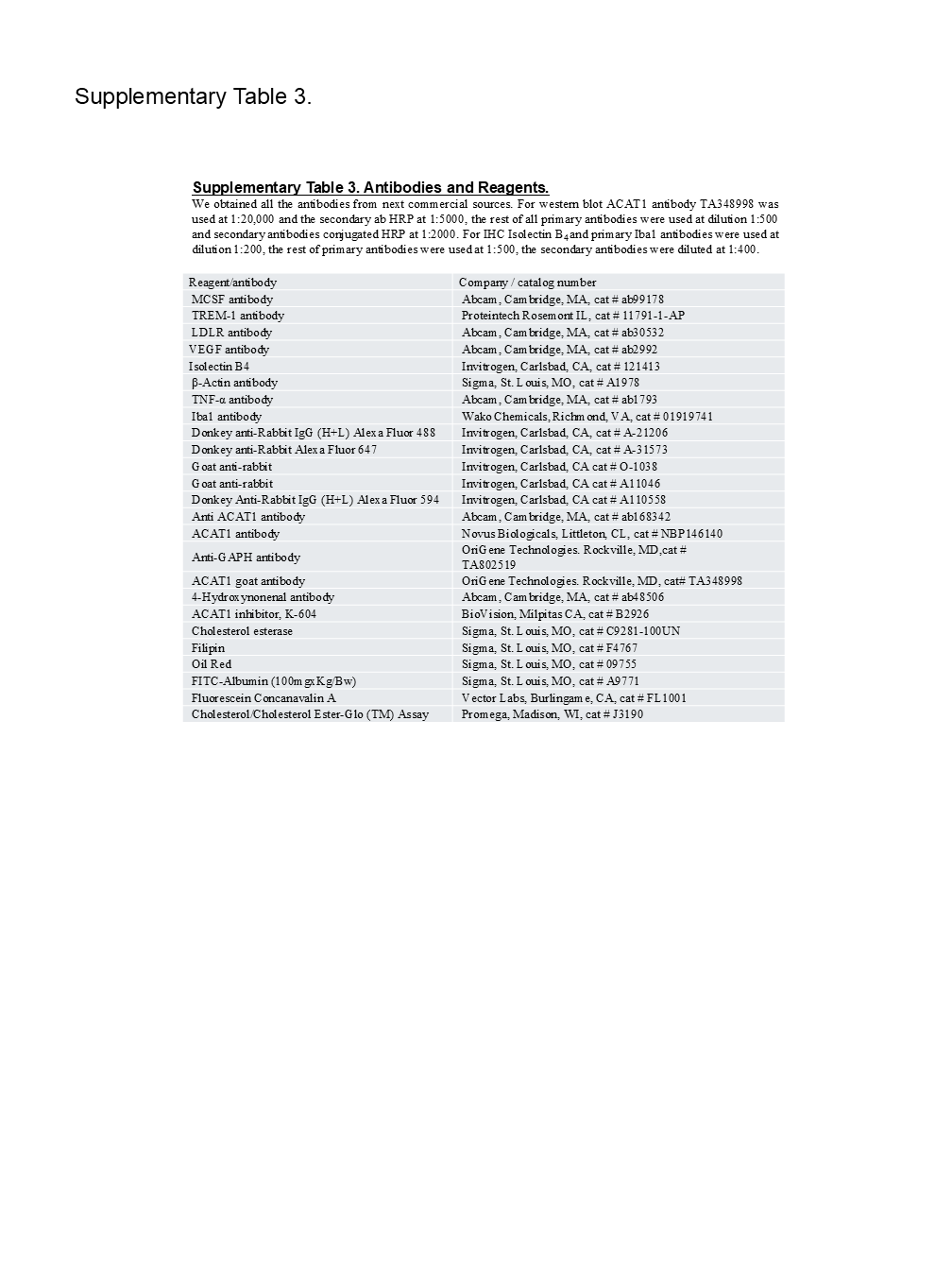

### Supplementary Table 4

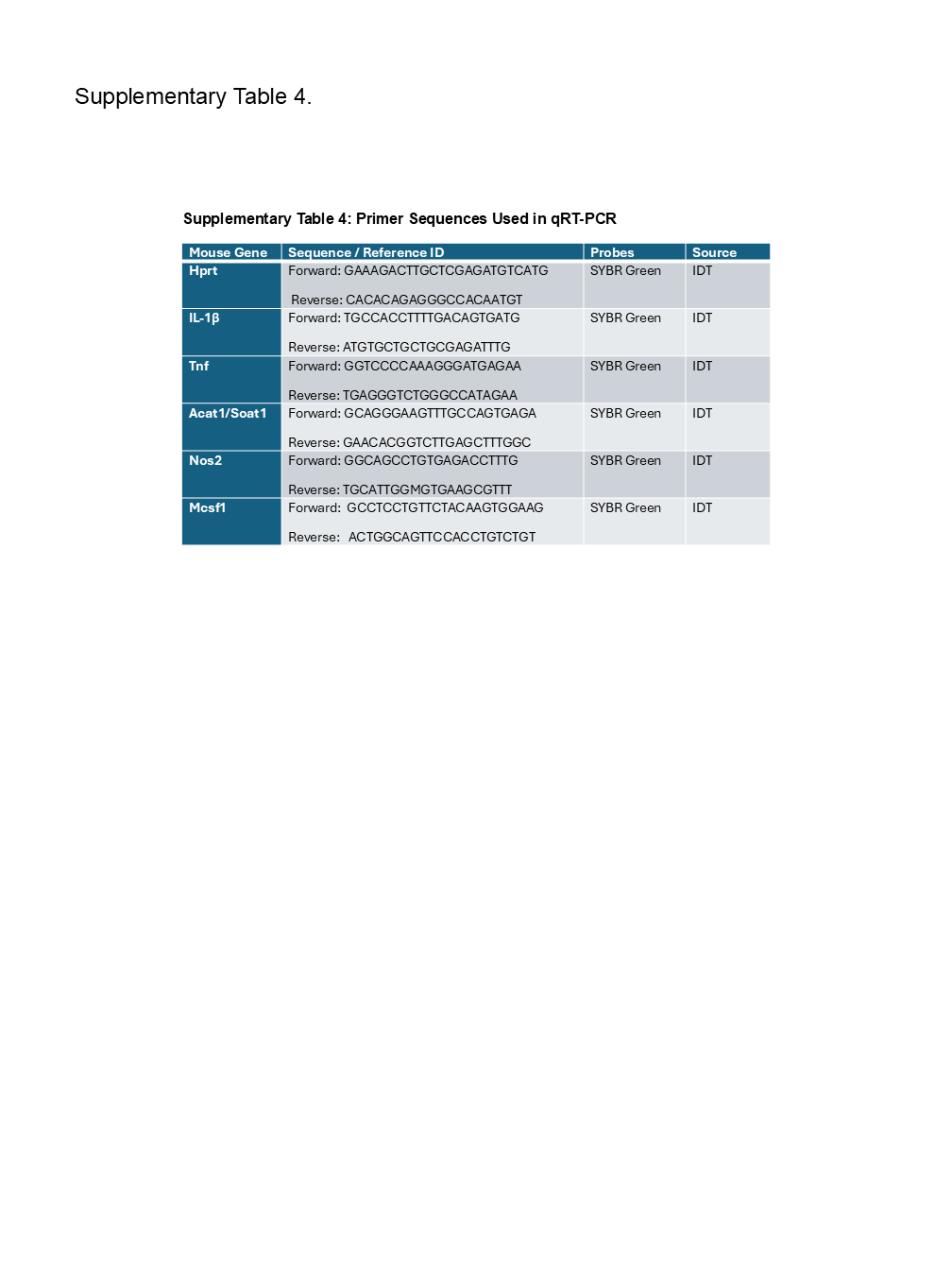
